# An improved Xer-cise technology for the generation of multiple unmarked mutants in Mycobacteria

**DOI:** 10.1101/758037

**Authors:** Yves-Marie Boudehen, Maximillian Wallat, Philippe Rousseau, Olivier Neyrolles, Claude Gutierrez

**Affiliations:** Institut de Pharmacologie et de Biologie Structurale (IPBS), Université de Toulouse, CNRS, UPS, 205 route de Narbonne, F-31400 Toulouse, France; Centre de Biologie Intégrative de Toulouse (CBI-Toulouse), Laboratoire de Microbiologie et de Génétique Moléculaires (LMGM), Université de Toulouse, CNRS, UPS, F-31400 Toulouse, France

**Keywords:** *Mycobacterium*, Xer-cise, *dif* site, recombineering, unmarked deletions, excisable cassette, mutagenesis

## Abstract

Xer-cise is a technique using antibiotic resistance cassettes flanked by *dif* sites allowing spontaneous and accurate excision from bacterial chromosomes with a high frequency through the action of the cellular recombinase XerCD. Here, we report a significant improvement of Xer-cise in Mycobacteria. Zeocin-resistance cassettes flanked by variants of the natural *Mycobacterium tuberculosis dif* site were constructed and shown to be effective tools to construct multiple unmarked mutations in *M. tuberculosis* and in the model species *Mycobacterium smegmatis*. The *dif* site variants harbor mutations in the central region and can therefore not recombine with the wild type or other variants, resulting in mutants of increased genetic stability. The herein described method should be generalizable to virtually any transformable bacterial species.

**Method summary:** *dif*-Zeo^R^-*dif* cassettes are used to replace non-essential genes in mycobacterial genome through recombineering. Spontaneous excision of the cassette is carried out under the action of the recombinase XerCD, resulting in unmarked deletions. Subsequent rounds of mutagenesis using cassettes flanked by a range of *dif* site variants allow construction of multiple mutants in which the different *dif* sites cannot recombine which each other, yielding stable genetic constructs.

## Main Text

Genetic studies of bacterial evolution and physiology require efficient and reliable tools for the construction of specific gene deletion mutants. A range of tools are available to manipulate Mycobacteria, such as *Mycobacterium tuberculosis*, the etiologic agent of tuberculosis. In particular, the development of phage recombinase-based recombineering, allowing replacement of a specific gene by an antibiotic resistance cassette through homologous recombination between linear DNA fragments and chromosomal DNA (Figure 1), has been a considerable improvement over previous methods [1]. However, only a limited number of selectable marker genes are available for mycobacteria [2] and, for safety reasons, strains used for medical studies must not carry resistance to multiple antibiotics. One way to overcome these problems is to use excisable cassettes that can be removed by recombination between two specific sites repeated at both sides of the cassette. The so-called Xer-cise technique is a convenient way to achieve auto-excision, because the recombination between two *dif* – for “deletion-induced filamentation” – sites is catalyzed by the XerCD recombinase present and active in most bacterial species as it is required for chromosome maintenance [3]. Xer-cise has been adapted to Mycobacteria, and shown to be efficient both in *M. tuberculosis* and in the fast-growing model species *Mycobacterium smegmatis* [2, 4, 5]. However, these studies used the wild type *dif* sequence of Mycobacteria [4, 6], leading to two possible drawbacks. First, XerCD-mediated recombination may occur between the *dif* sites on the linear allelic exchange substrate (AES) and the endogenous chromosomal *dif* site, which may generate false positive during the selection of antibiotic resistant recombinant clones. Second, after completion of the recombination events, the presence of several copies of the *dif* site on the bacterial chromosome may be deleterious because recombination between these sites may generate deletions or inversions on the chromosome, depending on the relative orientation of the *dif* sites [7]. Here we report an improvement of the Xer-cise technique in Mycobacteria. We designed and validated a set of variants of the natural *M. tuberculosis dif* site (Table 1) that can recombine with another copy of the same sequence, prompting excision of the antibiotic resistance cassette, but that recombine neither with the wild type nor with the other variant *dif* sites, yielding stable, unmarked multiple mutants.

**Table 1.** Sequence of wild type and variant *dif* sites^a^.

| *dif* site | Sequence |
| --- | --- |
| *M. smegmatis dif*<sup>+</sup> | AGTACCGATAA **GCTACA** TTATGTCAACT |
| *M. tuberculosis dif*<sup>+</sup> | TAAGCCGATAA **GCGACA** TTATGTCAAGT |
| *M. tuberculosis dif4* | TAAGCCGATAA **ATGATG** TTATGTCAAGT |
| *M. tuberculosis dif5* | TAAGCCGATAA **TAGAGT** TTATGTCAAGT |
| *M. tuberculosis dif6* | TAAGCCGATAA **CGGAAC** TTATGTCAAGT |
| *M. tuberculosis dif7* | TAAGCCGATAA **CTGATC** TTATGTCAAGT |

<sup>a</sup> left and right arm of the 28-bp *dif* sites are shown in black. The 6-bp central sequence is shown in red. Nucleotides modified in each variant are in bold letters.

**Figure 1.**
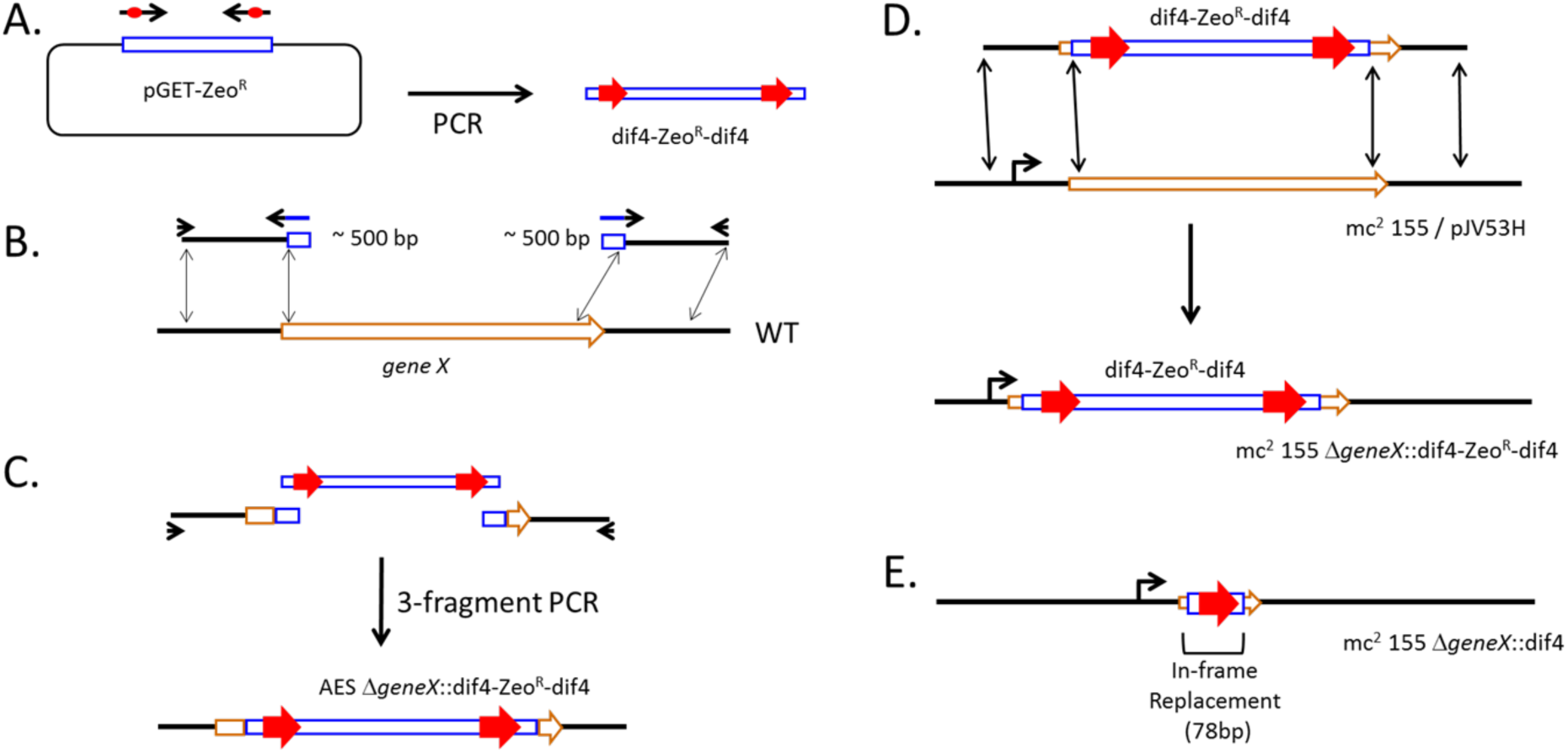
Construction of unmarked deletions in Mycobacteria by recombineering using zeocin-resistance cassettes flanked by *dif* site variants. **(A)** PCR amplification of Zeo^R^ cassettes with oligonucleotides introducing *dif* site variants on each side (red arrows). **(B)** PCR amplification of DNA fragments flanking the gene to be deleted. Blue boxes show sequence extensions identical to the left or right ends of the Zeo^R^ cassette. **(C)** 3-fragment PCR amplification yielding the allelic exchange substrates (AES). **(D)** Recombineering in Mycobacteria carrying plasmid pJV53H yields Zeo^R^ recombinant clones. **(E)** Spontaneous XerCD-dependent recombination between the tandem *dif* site variants generates unmarked deletion with a 78-bp scar replacing the deleted sequences.

### *dif* site variants

*dif* sites have been identified in *M. tuberculosis* and *M. smegmatis* [4, 6]. The minimal site is 28-bp long, with a XerC-binding left arm, a XerD-binding right arm and a 6-bp central region (Table 1). The mechanism of XerCD-mediated recombination has been well described in *Escherichia coli* [8, 9]. It involves the formation of a synaptic complex gathering two *dif* sites in a head-to-tail conformation inside a XerCD hetero-tetramer. The two XerD protomers create single-strand cleavage on each *dif* site, at the border of the central region, then promote strand transfer at the equivalent positions of the other site, leading to a Holliday junction-containing intermediate. By analogy with other site specific recombinase of the Y-recombinase family [8], efficient strand transfer requires that the first 2-3 nucleotides of the central region can pair. Indeed, recombination between sites with heterologous central region is inefficient [10, 11]. After isomerization of the intermediate, XerC protomers cleave both *dif* partners on the other side of the central region and transfer the second pair of strands by the same mechanism, allowing resolution of the intermediate and completion of the reaction. Both steps of reciprocal strand transfer require complementary sequences in the two recombination partners. Therefore, concerted modifications of the two nucleotides on both sides of the central region result in *dif* site variants still able to recombine with an identical site, because complementarity is conserved, but not with the wild type sequence with which base pairing at the border of the central region is no longer possible. Four variants, named *dif*4 to *dif*7 were designed following this rationale (Table 1). These variants cannot recombine with each other, allowing construction of stable strains carrying multiple variants in their genome. Using specific oligonucleotides (Table 2) and plasmid pGET-Zeo^R^ as a template, we PCR-amplified a set of Zeocin-resistance cassettes [12] flanked by the four variants of the *dif* site (Figure 1A).

**Table 2.**
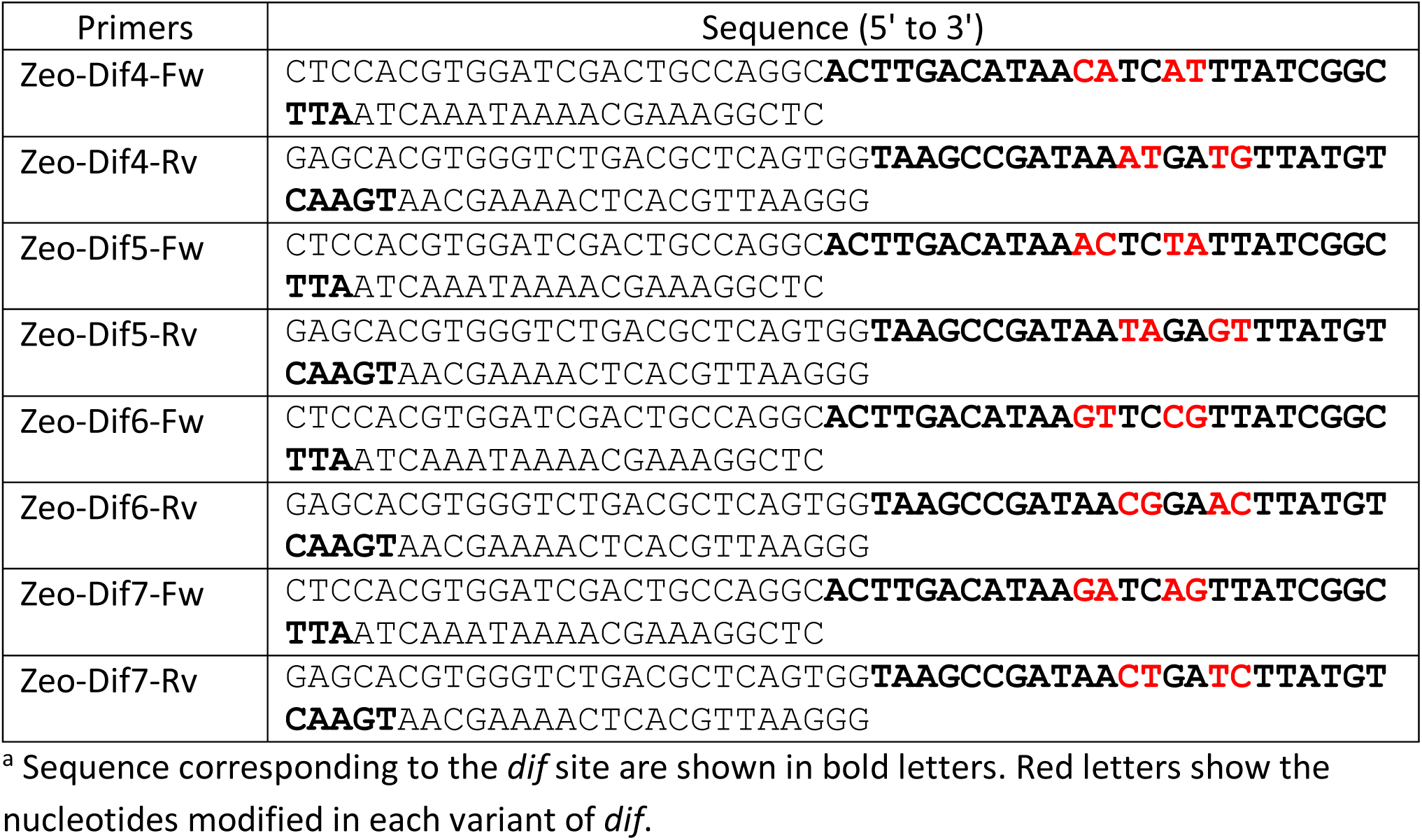
Oligonucleotides used to amplify Zeocin resistance cassette flanked by variant *dif* sites^a^

### Construction of unmarked deleted mutants

In order to delete a specific DNA segment, approximately 500-bp long upstream and downstream fragments were amplified by PCR using oligonucleotides that introduce sequences overlapping the right or left end of the *dif*-Zeo^R^-*dif* cassettes (Figure 1B, Table 2). Next, three-fragment PCR allowed amplification of an approximately 1,700-bp AES (Figure 1B-C). 100 ng of the purified AES were used to transform *M. tuberculosis* (strain H37Rv, ATCC27294) or *M. smegmatis* (strain mc^2^ 155, ATCC700084) carrying the plasmid pJV53H, a derivative of pJV53 [1], which expresses a mycobacteriophage recombinase and confers resistance to hygromycin (Figure 1D-E). Transformation was performed according to [1] and transformants were plated on Middlebrook 7H11 agar (Difco) supplemented with 10% oleic acid-albumin-dextrose-catalase (OADC, Difco) (complete 7H11 medium) and zeocin (25 µg ml^-1^; Thermo Fisher Scientific). Zeocin resistant (Zeo^R^) clones were re-streaked on the same medium and single Zeo^R^ colonies were inoculated in Middlebrook 7H9 medium (Difco) supplemented with 10% Albumin-dextrose-catalase (ADC, Difco) and 0.05% Tween 80 (complete 7H9 medium) without antibiotic for growth, DNA extraction and verification by PCR. Zeo^R^ clones exhibited replacement of the band corresponding to the wild type sequence by bands of the expected size for the *dif*-Zeo^R^-*dif* cassette insertion. These clones were streaked on complete 7H11 agar without antibiotic and grown at 37°C. Fifty single colonies were phenotypically tested in grids on complete 7H11 agar with or without zeocin, to screen for spontaneous excision of the cassette. Although at a variable frequency, we systematically observed Zeo^S^ clones among the 50 clones tested at this stage, irrespective of the *dif* variant of use, demonstrating that all of them were functional. Zeo^S^ clones were inoculated in complete 7H9 medium for growth, DNA extraction and PCR verification. For construction of multiple mutants, the verified Zeo^S^ Hyg^R^ clones were submitted to a subsequent round of mutagenesis, using a *dif*-Zeo^R^-*dif* cassette with a different *dif* site variant. Finally, clones verified to carry the expected mutation(s) were streaked on complete 7H11 agar without antibiotic and single colonies were phenotypically tested in grids on complete 7H11 agar with or without hygromycin, to screen for spontaneous loss of the pJV53H plasmid (Hyg^S^). The efficiency of this step could be improved by the utilization of a counter-selection marker or of a visual phenotype screening associated to plasmid pJV53H [5]. A final DNA extraction and PCR verification of each mutant was performed on the Zeo^S^ Hyg^S^ clones (see Figures 2 and 3).

**Figure 2.**
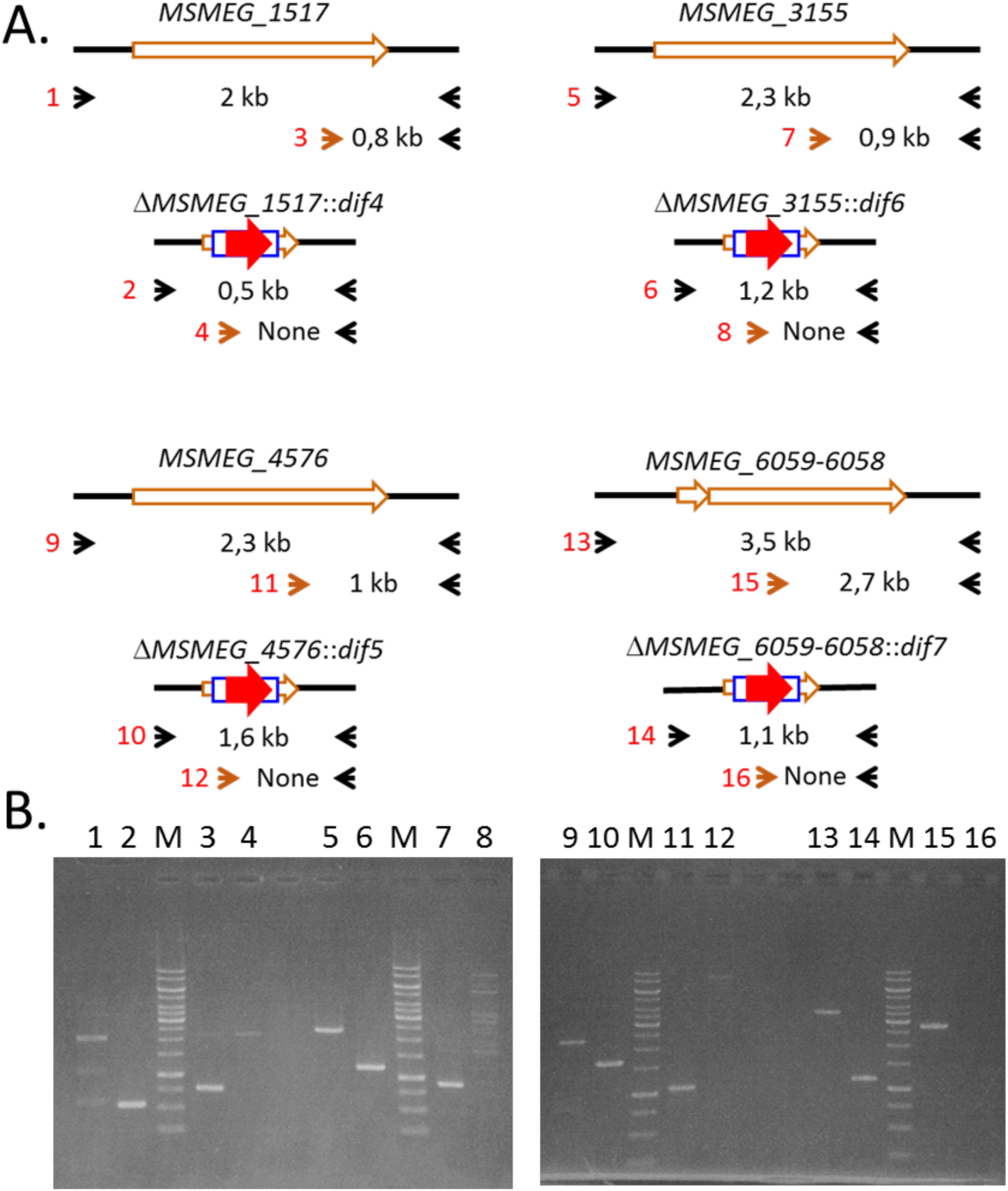
Verification of the quadruple mutant of *M. smegmatis* mc^2^ 155 ΔMSMEG_3155::*dif*6 ΔMSMEG_1517::*dif*4 ΔMSMEG_4576::*dif*5 ΔMSMEG_6059-6058::*dif*7. **(A)** Schematics of the primers (arrowheads) and PCR products used to verify the four deletions. Brown arrowheads indicate primers internal to the sequences to be deleted. Red numbers in front of each pair of primers refer to the lanes in 2B. **(B)** Agarose gel analysis of the amplified fragments. PCR were performed using DNA from wild type mc^2^ 155 (odd numbers) or the triple mutant strain (even numbers). M: 1 kb DNA ladder (Thermo Scientific).

**Figure 3.**
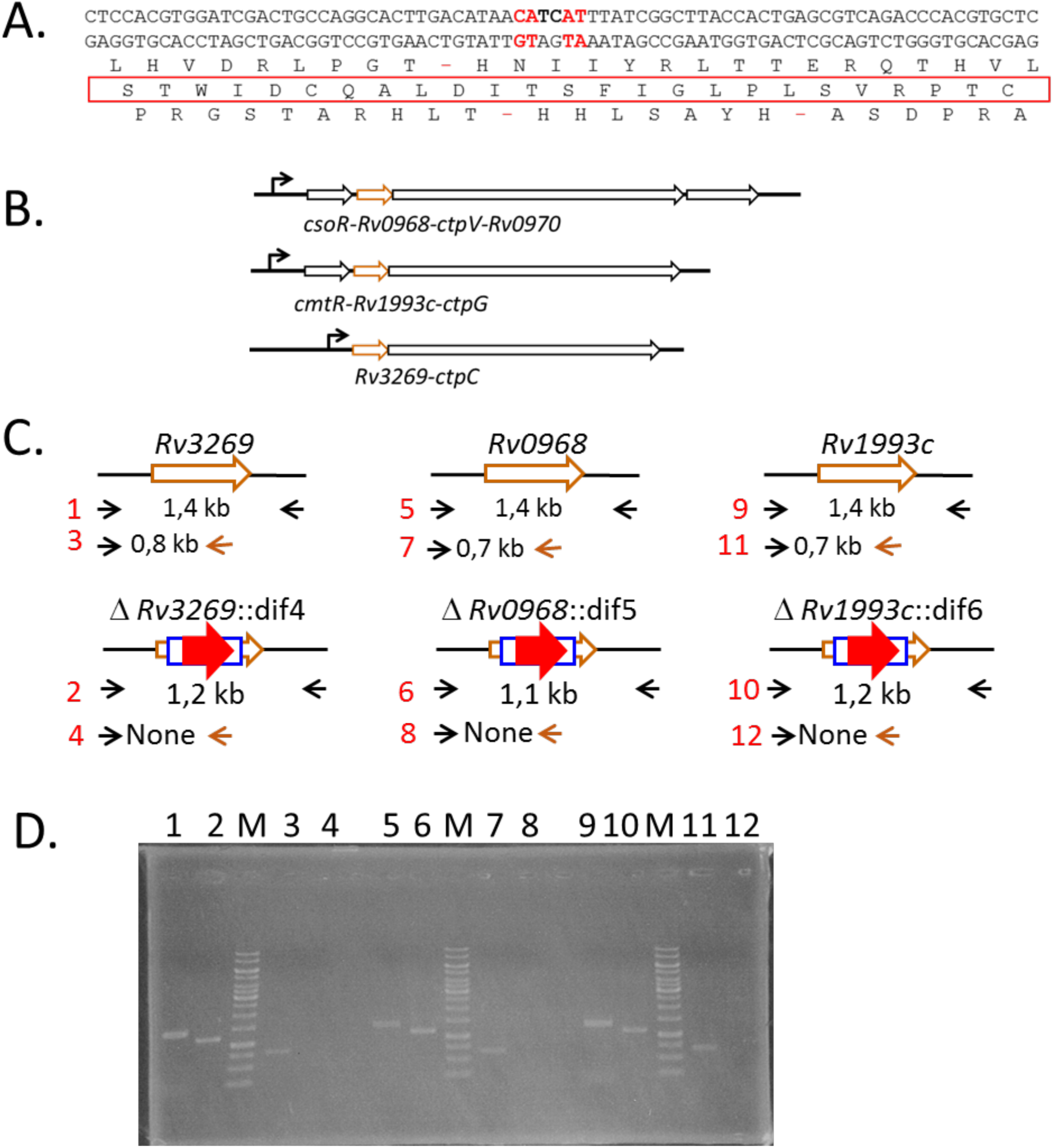
Construction and verification of H37Rv ΔRv0968::*dif*5 ΔRv1993c::*dif6* ΔRv3269::*dif*4, a triple in-frame deletion mutant of *M. tuberculosis*. **(A)** Translation of the 78-bp *dif*4-scar left after cassette excision **(B)** Genetic organization of the operons encoding *ctpC, ctpG* and *ctpV*. **(C)** Schematics of the primers (arrowheads) and PCR products used to verify the three deletions. Brown arrowheads indicate primers internal to the sequences to be deleted. Red numbers in front of each pair of primers refer to the lanes in 3D. **(D)** Agarose gel analysis of the amplified fragments. PCR were performed using DNA from wild type H37Rv (odd numbers) or the triple mutant strain (even numbers). M: 1 kb DNA ladder (Thermo Scientific).

### Construction of multiple unmarked deletions in *M. smegmatis*

To validate the functionality of this system, we constructed deletion mutants in *M. smegmatis* mc^2^ 155. Successive rounds of recombineering using AES constructed with appropriate primers (Table S1) and a different *dif* variant for each gene to be deleted allowed construction of multiple deletions. As an example, we constructed a quadruple mutant, namely mc^2^ 155 ΔMSMEG_1517::*dif*4 ΔMSMEG_4576::*dif*5 ΔMSMEG_3155::*dif*6 ΔMSMEG_6059-6058::*dif*7. DNA extracted from this mutant was amplified in PCR reactions with pairs of external primers or with an external primer and a second one internal to the deleted sequence (Figure 2A and Table S1). Analysis of the PCR products confirmed the genetic structure of the mutant (Figure 2B).

### Construction of multiple unmarked in-frame deletions in *M. tuberculosis*

Deletion of genes embedded within operons often results in polar effects on the downstream genes and an interesting advantage of excisable cassettes is the possibility to construct in-frame deletions, which circumvent such effect [4, 13]. After excision, the *dif*-Zeo^R^-*dif* cassettes used here leave a 78-bp scar with no stop codon in the +1 translation frame (Figure 1E and 3A). Therefore, provided that internal deletions of the target gene were designed in frame with this +1 frame, the final construction fuses the translation start to the 26 aa translated from this frame and the last codons of the deleted gene, which should not affect expression of the downstream genes. For example, the genes *ctpC, ctpG* and *ctpV*, encoding P-type ATPases involved in efflux of heavy metal ions in *M. tuberculosis* [14-16], are organized in operons containing an upstream gene encoding a small protein of unknown function, Rv3269, Rv1993c or Rv0968, respectively (Figure 3B). A triple mutant of *M. tuberculosis* deleted of Rv0968, Rv1993c and Rv3269, namely H37Rv ΔRv0968::*dif*5 ΔRv1993c::*dif*6 ΔRv3269::*dif*4, was constructed in three rounds of recombineering with appropriate AES (Table S2). The PCR verification of this triple mutant is shown in Figure 3D. Sequencing of the PCR fragments encompassing the variant *dif* sites confirmed the in-frame nature of the three deletions.

Finally, we believe that the improvement of the Xer-cise technique described here can be useful to construct mutants in all mycobacterial species prone to recombineering. Our approach is applicable to any antibiotic resistance cassette by adapting the primers used to amplify the cassette (Table 2). It can also be extended to the many bacterial species in which XerCD, or the functionally equivalent XerS or XerH systems are present, and to archaeal species in which a Xer-like system operates [17, 18].

## Author contributions

YMB, MW and CG performed experiments. ON acquired funding. CG and YMB wrote the manuscript; ON and PR edited the manuscript. All the authors contributed to conception and design, analysis and interpretation of data.

## Acknowledgements

We thank Vladimir Malaga and Stevie Jamet for the gift of plasmids pJV53H and pGET-Zeo^R^, respectively and François-Xavier Barre and François Cornet for helpful discussions.

## Financial & competing interests disclosure

This work was supported by Centre National de la Recherche Scientifique; Université Paul Sabatier; Agence Nationale de la Recherche (ANR14-CE14-0024, Fondation pour la Recherche Médicale (DEQ2016 0334902); Y.-M.B. is supported by FRM. The authors have no other relevant affiliations or financial involvement with any organization or entity with a financial interest in or financial conflict with the subject matter or materials discussed in the manuscript. No writing assistance was utilized in the production of this manuscript.

Papers of special note have been highlighted as: •• of considerable interest

## Supplemental material

**Supp Table S1.**
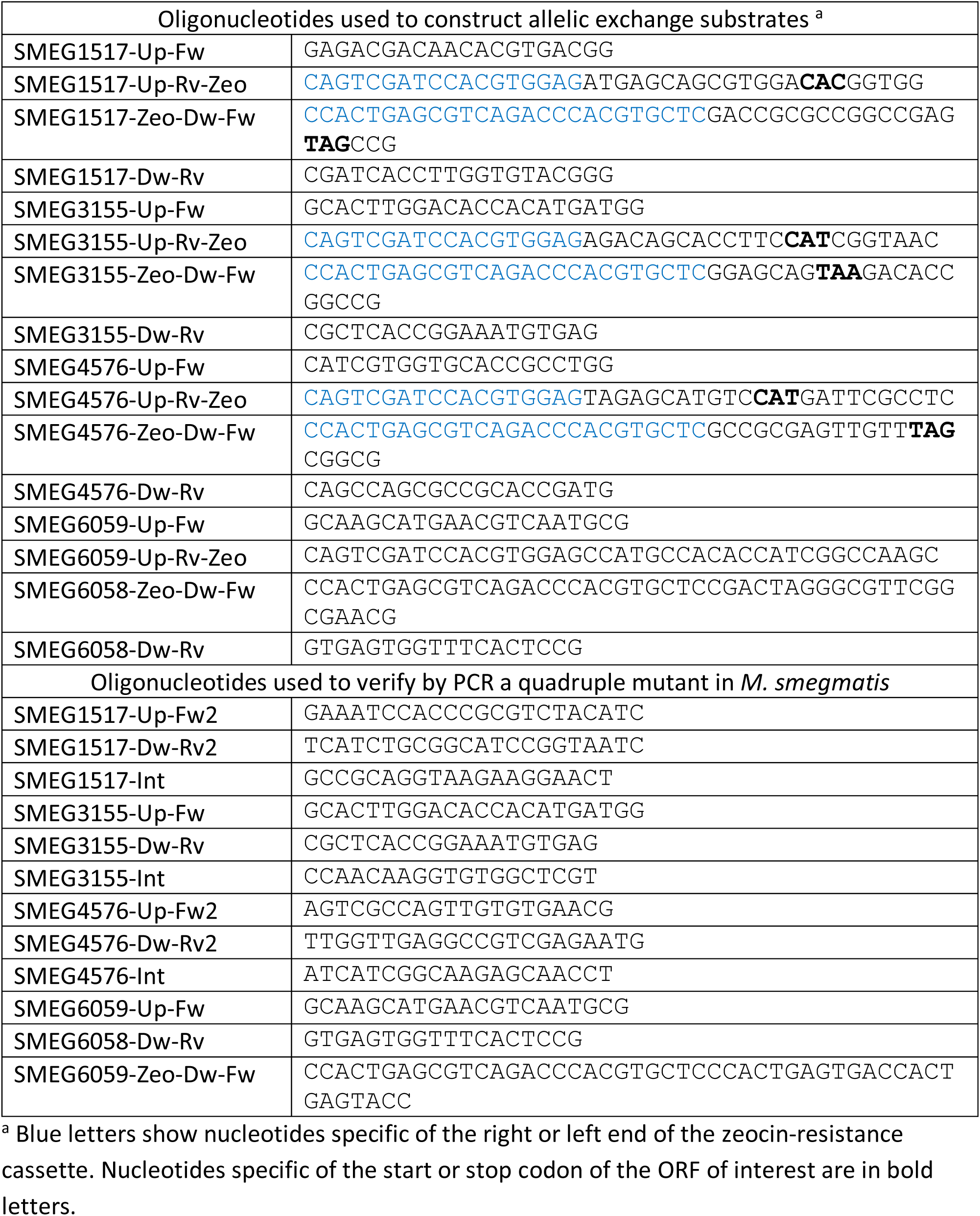
Oligonucleotides used to construct and verify a quadruple mutant in *M. smegmatis*

**Supp Table S2.**
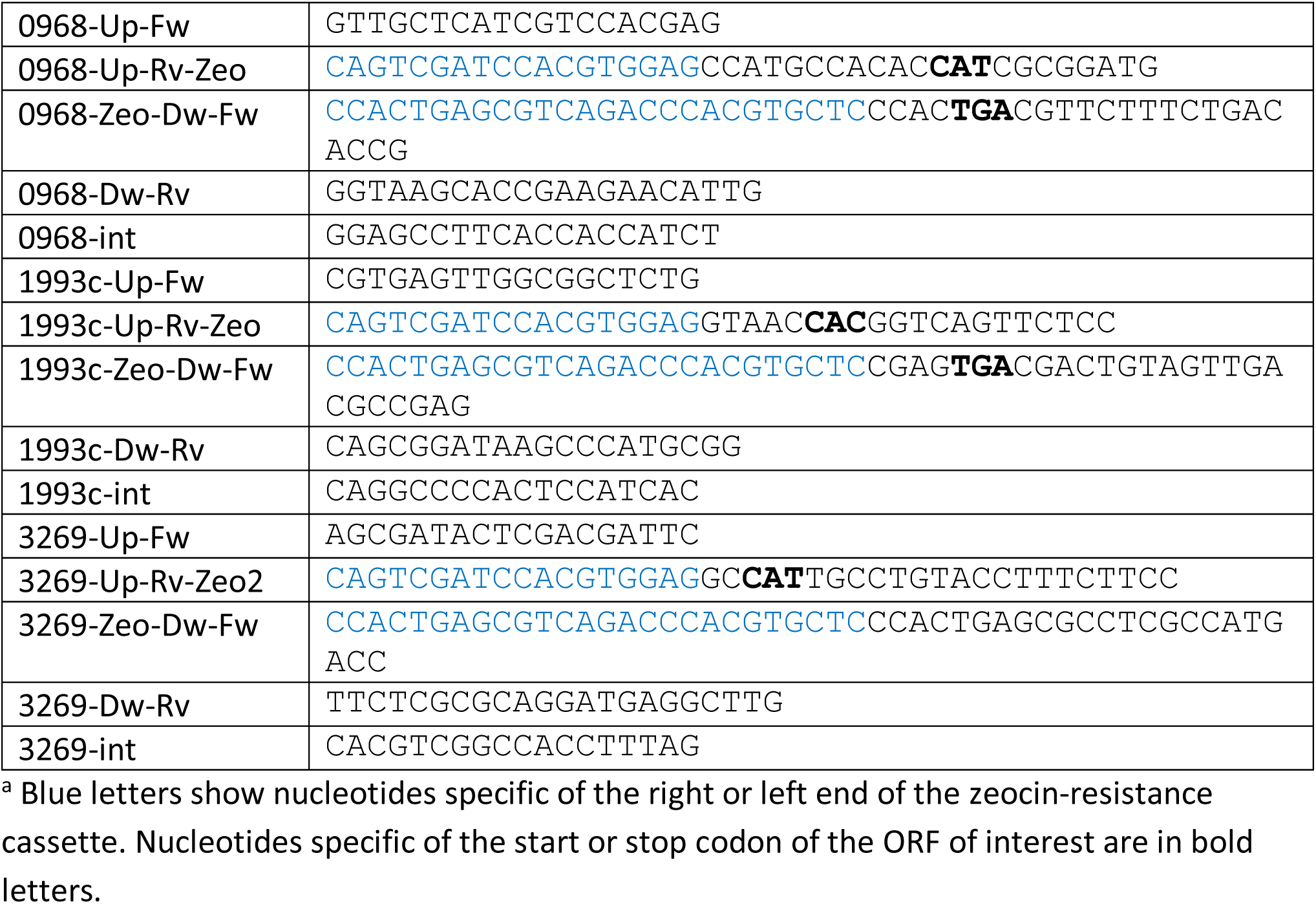
Oligonucleotides used to construct and verify a triple in-frame deletion mutant in *M. tuberculosis* ^a^

